# Intellectual Synthesis in Mentorship Determines Success in Academic Careers

**DOI:** 10.1101/273888

**Authors:** Jean F. Liénard, Titipat Achakulvisut, Daniel E. Acuna, Stephen V. David

**Affiliations:** Oregon Hearing Research Center, Oregon Health & Science University, Portland, Oregon, United States of America; Department of Bioengineering, University of Pennsylvania, Philadelphia, Pennsylvania, United States of America; School of Information Studies, Syracuse University, Syracuse, New York, United States of America

## Abstract

As academic careers become more competitive, junior scientists need to understand the value that mentorship brings to their success in academia. Previous research has found that, unsurprisingly, successful mentors tend to train successful students. But what characteristics of this relationship predict success, and how? We analyzed an open-access database of about 20,000 researchers who have undergone both graduate and postdoctoral training, compiled across several fields of biomedical science. Our results show that postdoctoral mentors were more instrumental to trainees’ success compared to graduate mentors. A trainee’s success in academia was predicted by the degree of intellectual synthesis with their mentors, resulting from fusing the influence of disparate advisors. This suggests that junior scientists should have increased chances of success by training with and linking the ideas of mentors from different fields. We discuss the implications of these results for choosing mentors and determining the duration of postdoctoral training.

## 1. Introduction

Most scientific researchers spend several years training under just one or two graduate and/or postdoctoral mentors, suggesting that this small number of relationships can have large impact on their subsequent career. Mentorship is believed to provide both direct intellectual benefits to the trainee – through the learning of new skills and concepts – and indirect social benefits – through engagement with the social network of the mentor ^28,42^. Reflecting this widespread sentiment, the stature of mentors and their letters of recommendation are given substantial weight in faculty hiring decisions ^17,32,50^. However, little is known about how the different stages of academic mentorship actually affect the protégé’s subsequent career ^36,49^. This question is not simply theoretical: identifying the individual determinants of academic success is urgent for trainees searching for faculty positions. More and more postdoctoral fellows are unable to secure a permanent research position even after years of additional training beyond their Ph.D. Trainees in this position must find ways to extend their postdoctoral training (“permadocs”) or join many of their colleagues in dropping out of academic research (the “postdocalypse” ^8,41^). Although this issue has gained attention recently, the plight of extended postdoctoral fellowships has been identified since it became a widespread practice, more than 35 years ago ^38^.

Success in academic research careers can be assessed by several different metrics, including publication and citation rates, funding levels, and a protégé’s own mentoring achievements. *Academic proliferation* (the number of progeny trained by a mentor, sometimes termed academic fecundity) provides a measure of this last metric ^15,33^. Empirical studies have found the number of academic progeny to be correlated with academic achievements such as holding a named chair ^14^, publishing more papers ^34^ or receiving the prestigious Nobel prize ^7^. Academic proliferation provides a measure of two factors: (1) attrition rate, where a researcher who has never mentored someone else probably does not hold a permanent position, and (2) scientific proficiency, where more successful mentors have a greater number of trainees. This second effect might reflect that greater fame attracts more students, greater financial resources allows more hires, and a virtuous circle where trainees contribute back to the prestige of the mentor through collaboration and contribution to an extended social network throughout their own careers ^42^.

Given the central role that mentorship plays in academic research, studying a large network of mentors and trainees has the potential to provide insight into the drivers of academic success. The Academic Family Tree (*academictree.org*) is an online effort begun in January 2005 to document training relationships in a relational database. This project originally started with a focus on the field of neuroscience but progressively expanded to span more than 30 disciplines ^15^. Researchers in the database are linked to publications they have authored by automated record linkage to the Medline database. In the current study, we applied a data-driven approach to study 500,000+ life science researchers, with a focus on the subset with documented graduate and postdoctoral training. Our objective was to uncover how patterns in the network of mentors and protégés stage their academic success, with a focus on several questions: To what extent does mentorship impact the future career of trainees? What is the relative influence of social versus intellectual relationships mentoring relationships? Do graduate or postdoctoral mentors have a greater impact on trainee careers? What are the long-term temporal trends underlying the success of trainees?

To address these questions, we identified several properties of the mentor network graph and of semantic relationships between publications by mentors and trainees. We then used a regression framework to quantify the impact of these different factors on the two outcome variables defined above: acquiring an independent research position after postdoctoral training and the academic proliferation of those who do obtain independent positions. Our analysis revealed that several factors had significant predictive power for the success of independent careers. Trainees of graduate and postdoctoral mentors with high proliferation tended to be more successful themselves, consistent with the previous observations for graduate mentors ^34^. In addition, success rates were predicted by the pattern of intellectual similarity between mentors and trainees. Trainees whose research was able to synthesize the influence of intellectualy disparate mentors had larger odds of continuing in research. Thus a model emerges in which the most successful trainees are trained by highly connected mentors and are able to synthesize content of both mentors’ work in their own research.

## 2. Results

### 2.1 Properties of mentorship networks

We evaluated several population-level features of triplets in life science fields documented in the Academic Family Tree. These features group broadly into two categories: graphical properties of the mentorship network and semantic properties of mentor and trainee publications (Fig. 1). The first group includes the date of graduation and duration of postdoctoral training, proliferation rate of mentors and trainee (average number of trainees per decade), professional age of mentors (years since the mentor has completed training) and the distance completed training) and the distance to the mentors earliest common ancestor (Fig. 1A). The second group includes the similarity of scientific output by researchers in the triplet, measured by latent semantic analysis of published abstracts (Fig. 1B-C). The specifications of these variables are detailed in the Methods (Section 5.5).

**Figure 1:**
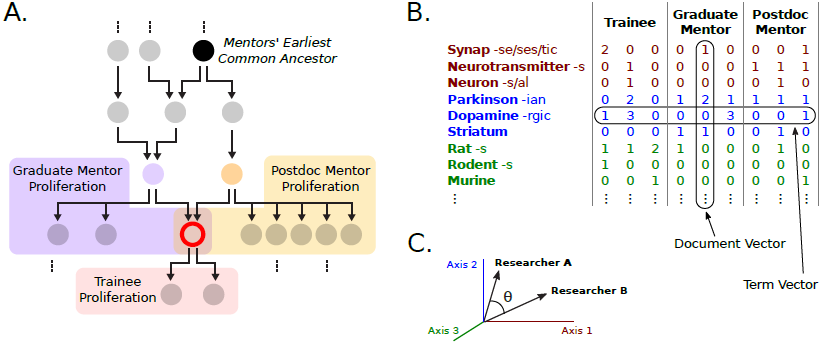
Predictor variables. (A) Schematic mentorship network graph for a single trainee (red), with mentor-trainee relationships indicated by arrows. Analysis focused on researchers who completed training with at least one graduate (purple) and one postdoctoral mentor (yellow). Mentor network distance is computed as the distance to their earliest common ancestor (black). Professional success is measured by proliferation rate, the average number of trainees per decade after the start of a researcher’s independent career. (B) Intellectual relationships were characterized by semantic analysis of abstracts indexed in the Medline and Scopus publication databases. Semantic content was quantified by a term frequency–inverse document frequency (TF-IDF) metric, in which stemmed words were counted and their relative frequency was used to define a space for principal component analysis. (C) The term vector for each publication abstract was projected onto a vector spanning the 400 largest principal components. Publication similarity was computed as the cosine distance between the average publication vector for each researcher, after excluding their commonly authored publications.

The primary measure used to evaluate success of academic careers of both trainees and mentors is the academic proliferation rate, defined as the average number of researchers trained per decade. Across all triplets, both trainees and their mentors show overall similar distributions of proliferation rates, with a mode of about one (*i.e.* one trainee every 10 years, Fig. 2A). Mentor proliferation rates are well-described by a Poisson distribution. However, 60% of the protégés did not train anyone themselves, reflected by a large peak at 0 proliferation rate. Their distribution is better described by the product of binomial and Poisson distributions (see below and Eq. 1). The proliferation rates of graduate and postdoc co-mentors within a triplet have low correlation (Pearson’s coefficient r = 0.14 with 95% CI [0.12, 0.16]), producing a wide distribution of the difference in proliferation rates (Fig. 2B, blue bars). This distribution is not symmetric. Postdoctoral mentors had significantly greater average proliferation than graduate mentors (mean difference: 4.50, 95% CI [4.24, 4.75]). Although their proliferation rates differed, co-mentors tended to be closer in the training network than expected by chance (Fig. 2C). Protégés in triplets were trained by mentors with an average academic age of 10 years (Fig. 2D). Graduate mentors tended to have slightly lower academic age than postdoc mentors, possibly reflecting budget constraints on hiring more costly postdocs during the early independent career. This age difference is stable through time (data not shown).

**Figure 2:**
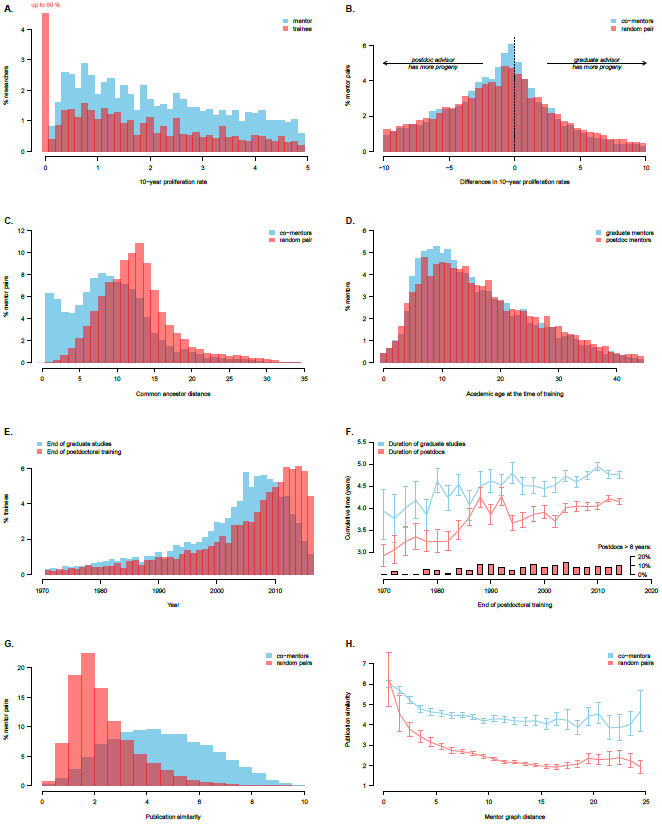
Main features of the *n* = 18, 856 mentorship triplets in life science analyzed in this study. Each triplet is constituted by a pair of graduate and postdoc mentors and their common trainee. A: distribution of proliferation rate (average number of trainees per decade) of mentors (blue) and trainees (red). The large peak at zero for trainees reflects the large number of trainees that do not go on to mentor students of their own. B: difference in proliferation rate between graduate and postdoc mentors common to triplets (blue) and mentor pairs picked at random (red). C: common ancestor distance of co-mentors (blue) and mentors picked at random. D: academic age at the time of training for graduate and postdoc mentors of each triplet. E: year of graduation (blue) and end of postdoc (red) for triplets. The decreasing number of trainees graduated in recent years, in blue, results from the inclusion criterion of trainees that were selected as those having undergone postdoctoral training. F: Mean cumulative time spent in graduate studies (blue) 6 and postdoctoral fellowships (red), as a function of postdoc end date. Postdoctoral studies lasting more than 8 years were excluded from the computation of cumulative time. Instead their proportion was plotted as bars at the bottom the graph. G: the similarity between co-mentors (blue) is higher than among a randomly picked pair of researchers (red). H: closer common ancestor distance leads to greater publication similarity, and this effect is cumulative with the higher proximity of researcher that co-mentor the same trainee.

Some long-term trends are readily visible in the data set. Most triplets in the analyzed data completed training after 1990 (Fig. 2E). This upward trend reflects the ongoing growth of postdocs in life science ^39^, the number of which increased by a factor of four over the period 1980-2010 ^49^. The ten-fold increase reported in Fig. 2E is larger than this trend. This difference may reflect the substantial recent growth of the well-represented field of neuroscience (^2^, Fig. 7), but it may also reflect a sampling bias in the database favoring more recent graduate students and postdocs. In parallel with the growing number of postdocs, the data also indicate an increase in the duration of training over time (Fig. 2F). This increase is true for the duration of both graduate and postdoctoral training, the latter of which has increased by an average of about about a year since the 1970s.

Research performed as a graduate student or postdoc may be more or less aligned to the mentor’s own research. To study how the similarity of intellectual output between mentors and their trainees impacts the subsequent success of trainees, we performed a latent semantic analysis on the abstracts of non-coauthored papers published before the end of postdoctoral training (see Methods and Fig. 1). Not surprisingly, co-mentors tended to have more similar semantic content in their publications than randomly selected pairs of researchers (Fig. 2G). Their publication similarity increased with proximity in the training network, as measured by mentor graph distance (Fig. 2H). However, graph distance is not the only factor influencing the publication similarity. Co-mentors displayed a greater similarity than randomly chosen pairs of researchers with the same graph distance (Fig. 2H). Thus, the relatively high publication similarity of co-mentors reflects factors beyond the similarity of their academic genealogy.

### 2.2 Intellectual synthesis is linked to continuation in academic research

Research careers adopt many different shapes and may involve a variable mix of research and university-level teaching. This seldom occurs outside of academia, in the private sector. Here, we focus specifically on a criterion that is a good proxy characterizing the success of academic research careers in life science, and which is accessible in our dataset: the training of at least one graduate student or postdoc fellow. Indeed, the training of a more junior researcher is a years-long commitment, and a stable research position is often an institutional prerequisite for it. Conversely, having postdocs and graduate students is the norm in life science research, and virtually all successful researchers in life science manage a team composed of graduate students and postdocs.

To investigate the impact of publication similarity on trainee academic success, we considered its relationship to the odds of becoming a mentor, *i.e.*, for the trainee to continue in academia and themselves train at least graduate student or postdoctoral fellow. We observed several significant correlations between publication similarity and trainee success. In particular, greater publication similarity between a trainee and each mentor led to higher probability of continuing in academia (Fig. 3A-B, *p<0.001*, two-sample Kolmogorov-Smirnov test). In contrast, co-mentors with greater publication similarity had trainees *less* likely to continue in research (Fig. **3C**, *p<0.001*). Together, these observations are consistent with a model in which a trainee who successfully synthesizes knowledge and approaches from dissimilar mentors is more likely to continue on to an independent academic career. Furthermore, successful trainees tended to show closer semantic proximity to the postdoctoral mentor than the graduate mentor (Fig. **3D**, *p<0.001*), suggesting that the postdoctoral relationship is a stronger determinant for the trainee’s future employment.

### 2.3 Model of academic success in life science

The patterns in Figs. 2 and 3 suggest a link between the characteristics of mentors and their trainee’s odds of staying in academia. At the same time, the relatively strong coupling between variables such as postdoc duration and training end date (Fig. **2F**) and mentor graph distance and publication similarity (Fig. **2H**) presents a challenge for building the predictive model of trainee success. In order to disentangle these factors, we developed a statistical model to measure the impact of each variables on the subsequent career of trainees. Our model considers two possible scenarios (Eq. 1): (a) the protégé moves on to a private sector position or to a public research position that does not involve training, thus excluding him from having descendants in the training network; and (b) the protégémoves on to an independent research position involving training, in which case their own proliferation rate is used to measure their human-capitalized scientific legacy. In order to permit adequate time for measuring trainee proliferation, we restricted our dataset to triplets where the protégéfinished their latest training no later than 2007. All the variables were available at the completion of postdoctoral training and thus were not biased by the subsequent independent career of protégés.

**Figure 3:**
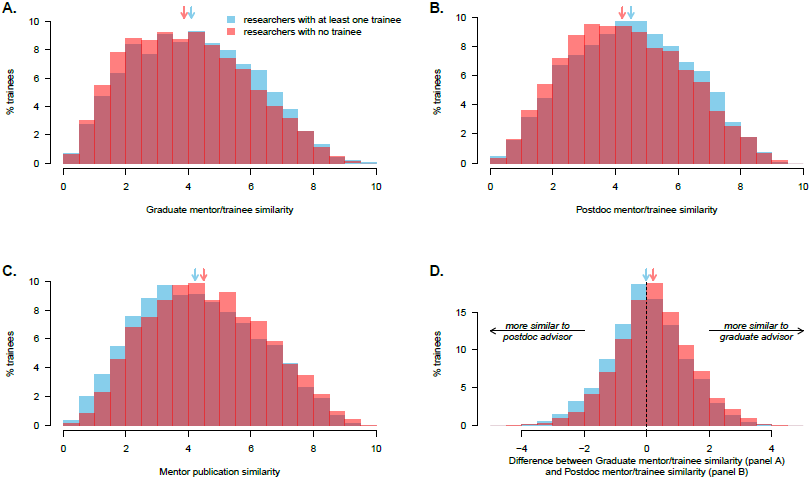
The odds of becoming an academic mentor are correlated with the trainee’s ability to synthesize disparate influences from mentors, as revealed by the similarity of their abstracts published prior to the end of the postdoc. Arrows indicate the medians (the differences are significant in all panels, *p<0.001*). A-B: Trainees who become mentors showed greater similarity with their graduate (A) and postgraduate (B) advisors. C: Lower similarity between mentor is linked with better odds to continue in academic research. D: Protégés that have a greater publication proximity with their postdoc mentors, compared to their graduate mentors, tend to move more often to independent academic positions.

These outcomes are integrated into a mixed model. Two model composition techniques are suitable to handle the large prevalence of postdocs without trainees, the *hurdle* and *zero-inflated* frameworks ^6^. They differ by their modeling of researchers without trainees: the hurdle framework assumes that all independent researchers have at least one trainee, while the zero-inflated framework allows the existence of some independent researchers that have no trainee in the database. This latter scenario would correspond either to incomplete Academic Tree profiles or to researchers not involved in graduate/postdoctoral training. Furthermore, the proliferation rate may be modeled as a Poisson distribution or as a negative binomial distribution. The former assumes that count variance is directly proportional to mean count, while the latter relaxes this assumption and allows over-dispersion, at the cost of an extra free parameter.

To decide which of the four model architectures (hurdle or zero-inflated with Poisson or negative binomial distribution) and which predictors yielded the best fit to the data, we screened the performance of each architecture on all possible combinations of predictors. This was done by calculating parameter values for each model that maximized log-likelihood of predicted outcomes, using a 10-fold cross-validation scheme. Shapley values were then computed scoring the relative contribution of each variable to the overall model’s performance. Negative binomial distributed count rates consistently outperformed count rates conforming to a Poisson distribution, indicating the presence of over-dispersion in rates for the highest training researchers. Zero-inflated models performed slightly better than hurdle models, indicating that an assumption that all permanent research faculty must have at least one trainee is not consistent with this dataset (Table 3). The best-fitting model was overall the zero-inflated negative binomial model, whose cross-validated predictions are shown along each dimension in Fig. 9 in Supplementary Information. We focus on this specific mathematical model for the remainder of the paper.

**Figure 4:**
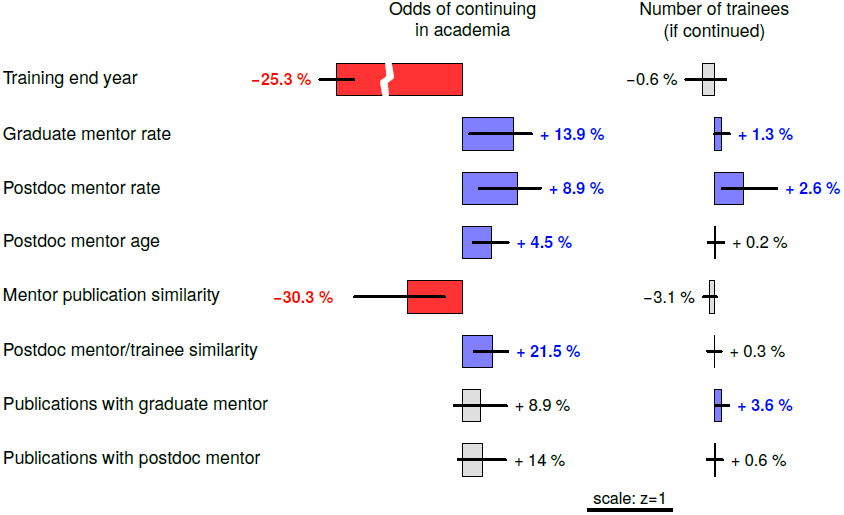
Modeled effects of variables for continuing in academia (left column) and for training rate when continuing (right column). The probabilities depict the change in the odds of continuing in academia (left) and in the training rate (right) induced by an increase of one unit for each variable. The error bars show the 95% bootstrapped confidence interval. Width of the bars and error bars show the relative importance (z-scores, see scale bar) whereas the percentages show the effect of adding one original unit. Significant changes are color-coded and the associated effects are shown in bold. For example, the interpretation for the “Postdoc mentor rate” variable is as follow: *ceteris paribus*, an increase of 1-point on the postdoc mentor proliferation scale (i.e. one more trainee per decade) results in increased odds by 9.3% of the protégés to find a permanent position, and this effect is statistically significant. The long term effect this change is also significant, and the protégé’s proliferation rate is then increased by 2.6%.

**Figure 5:**
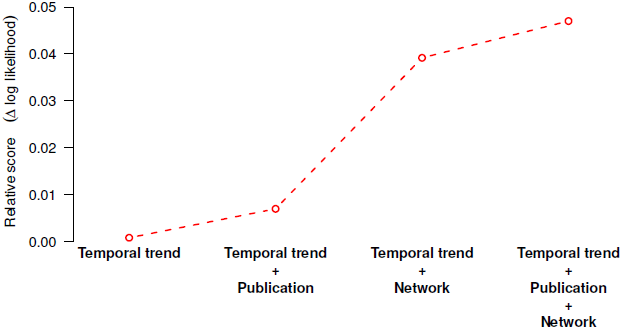
Impact of adding the different groups of variables on the log-likelihood of the model. Once the temporal trends are accounted for, the largest effect corresponds to the inclusion of mentor network features (training rate and academic age).

**Figure 6:**
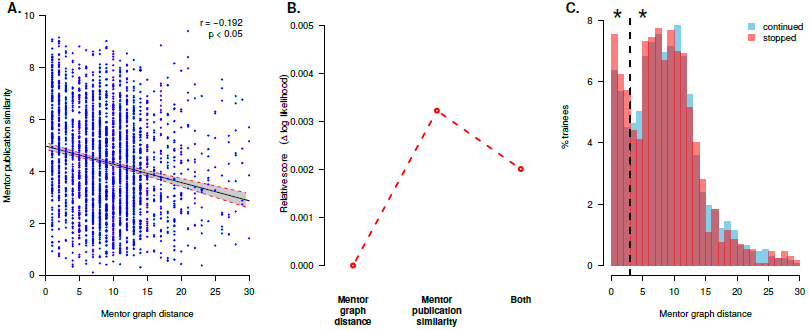
Relationship between the mentor early similarity in publications and their common ancestor distance, and their respective predictive power in the model. A: scatter plot with linear regression, the shaded area displays the 95% confidence interval in prediction. B: Relative contribution to the model goodness-of-fit, when acting in isolation or together. C: distribution of Common Ancestor Distance for the trainees who stopped (red) or continued (blue) a research academic career. The dashed line indicates the boundary between the two peaks in the distribution of mentor graph distance. Probability of continuing is significantly smaller for the group with shorter graph distance.

**Figure 7:**
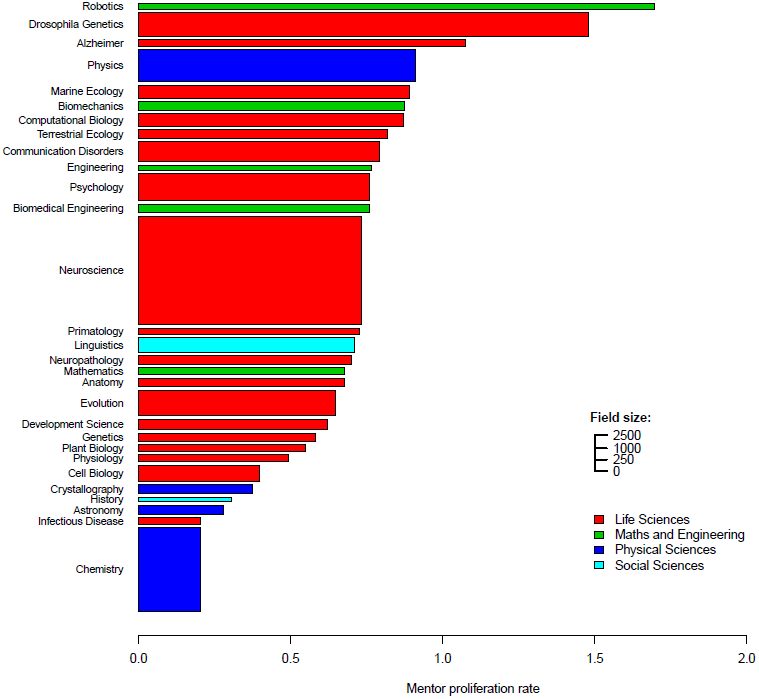
Statistical mode of the mentor proliferation rates (horizontal axis) and sample size (number of triplets along the vertical vertical axis). Life Sciences (red) are well represented in the Academic Family Tree dataset, and thus were the focus of the current study. The proliferation rate is between 0.05 (one trainee every 20 years) and 0.1 (one trainee every 10 years) in most of its sub-fields. Maths and Engineering fields (green) show generally higher proliferation rates, while Physical Sciences (blue) and Social Sciences (cyan) show lower rates.

**Figure 8:**
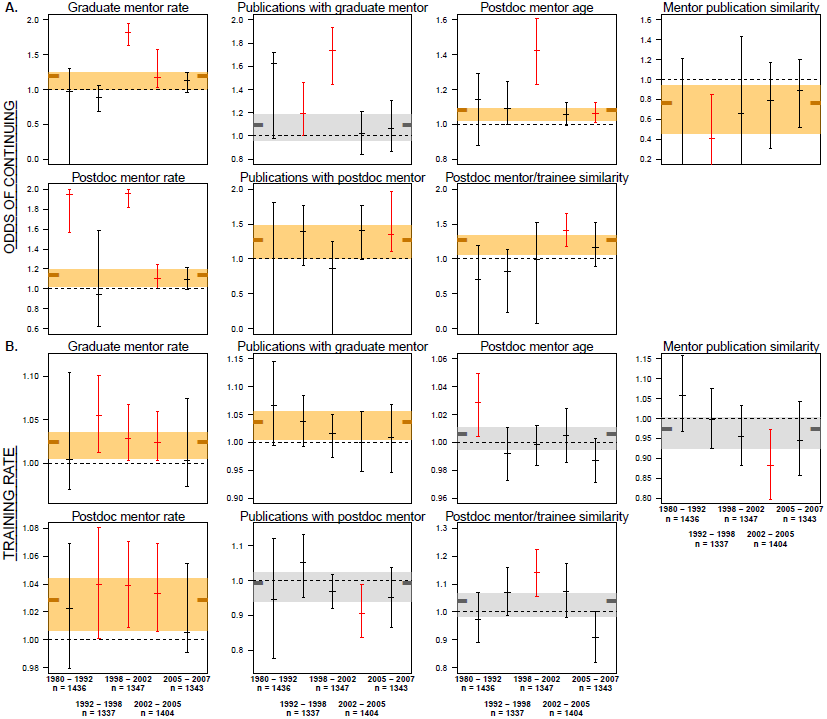
Regression coefficients obtained when optimizing the model on different temporal subsets while removing the postdoc end date from the set of predictors. Temporal partitions were chosen to contain roughly equal sample sizes, and were as follow: 1980 – 1992 (*n* = 1, 436); 1992 – 1998 (*n* = 1, 337); 1998 –2002 (*n* = 1, 347); 2002 – 2005 (*n* = 1, 404); 2005 – 2007 (*n* = 1, 343). The coefficients estimated to predict continuation in academia (A) and long-term mentoring rate (B) are consistent with the ones obtained with the full model that includes temporal predictors (represented in these plots as shaded areas). Specifically, whenever a variable reaches significance within a temporal subset, it does so in the same direction as the trend apparent in the full model. Reciprocally, whenever a variable reaches significance in the full model, there is at least one temporal subset when it reaches significance as well.

**Figure 9:**
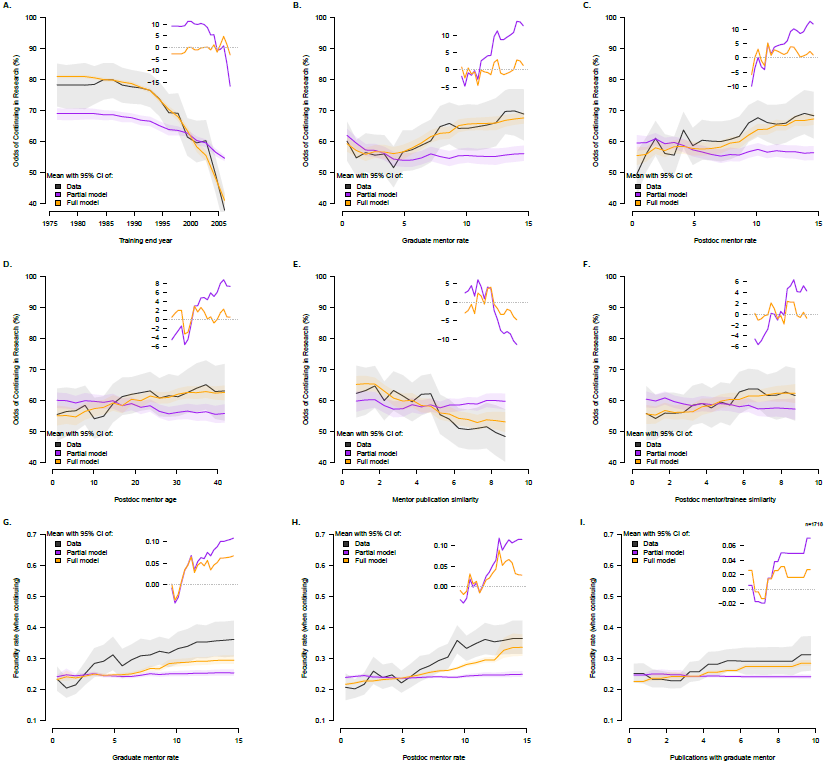
Academic continuation in life science (panels A to F) and modeled training rate (panels G to I) explained by the most significant variables. For each variable, the data is shown in black, and the cross validated predictions of the model including this variable is shown in orange (“full model”). To visualize the contribution of each variable, we also display the cross-validated predictions of the model without this variable in purple (“partial model”). Lines and shaded areas represent respectively the mean values and their 95% bootstrapped confidence interval.

Several variables had a negative Shapley score (Table 1), indicating model over-fitting. These variables tend to form spurious relationships with the protégé’s proliferation, limiting their relevance for modeling. In particular, three variables exhibited high levels of over-fitting: mentor protégénetwork distance, the academic age of graduate mentor, and the total time spent by the protégé’ in postdoctoral studies. These same variables were also systematically excluded by the iterative variable selection processes based on Shapley values (both CSA forward and backward algorithms, Table 1), thus we excluded them from the modeling. The relevance of the number of publications number with postdoc mentor is more ambiguous: it has a negative Shapley value, but is ranked above the number of publications with graduate mentor in the forward CSA algorithm. This metric is widely used to evaluate job applicants (for example, it has been reported as the most important metric used by committees studied in ^46^, ranking above the quality of journals or the funding track-record), thus we opted to include it in the model.

**Table 1:** Overview of factors impacting trainee proliferation and their contribution to the model’s goodness-of-fit. Highlighted variables are the ones kept in the model. The contribution of each variable to the overall fitness is Shapley value, with positive values denoting better fit. The 12 variables are ranked according to their increasing order of importance, according to the rank of their Shapley Value (“SV”), the rank when using the Forward Selection Algorithm (“FSA”) and the rank when using the Backward Selection Algorithm (“BSA”).

|  | Goodness-of-fit | Variable ranks |  |  |
| --- | --- | --- | --- | --- |
|  |  | SV | FSA | BSA |
| **Temporal trend** |  |  |  |  |
| Training end year | 1.822 | 1 | 1 | 2 |
| Postdoc duration | -0.027 | 10 | 11 | 11 |
| **Network** |  |  |  |  |
| Graduate mentor proliferation | 1.301 | 3 | 3 | 3 |
| Postdoc mentor proliferation | 1.764 | 2 | 2 | 1 |
| Mentor graph distance | -0.109 | 12 | 12 | 12 |
| Graduate mentor age | -0.010 | 8 | 7 | 8 |
| Postdoc mentor age | 0.121 | 6 | 6 | 7 |
| **Publication** |  |  |  |  |
| Mentors publication similarity | 0.369 | 4 | 4 | 5 |
| Graduate mentor/trainee similarity | -0.025 | 9 | 10 | 10 |
| Postdoc mentor/trainee similarity | 0.163 | 5 | 5 | 4 |
| Publications with graduate mentor | 0.090 | 11 | 9 | 6 |
| Publications with postdoc mentor | -0.004 | 7 | 8 | 9 |

### 2.4 Determinants of academic success

The impact of variables on the odds of obtaining of an independent research position and on long-term training rates are summarized in Table 1 and Fig. 4. The odds of continuing in academia were positively influenced by higher mentor proliferation rates, greater postdoctoral mentor academic age, and close publication similarity between the protégés and their postdoctoral mentor (Figs. 4 and 9B,C). In contrast, training end year s and high mentor publication similarity negatively influenced the probability of continuing in academia (Figs. 4 and 9A,D).

A similar, but not entirely overlapping set of variables influenced the protégé’s long-term proliferation rate. Trainee proliferation was positively influenced by the mentors’ proliferation rates, along with semantic proximity to the postdoctoral mentor and the number of publications co-authored with the graduate mentor (Figs. 4 and 9G-I). Overall, these results show that highly prolific mentors tend to provide their protégés with the assets required for their own success in academia, both in terms of securing permanent research positions and of increasing their long-term proliferation.

The effect of training end date on the odds of continuing in academia was found to be very strong (Figs. 4 and 9A), consistent with known trends toward a shortage of independent academic positions available to postdoctoral trainees (e.g. ^26,37,38^). We considered the possibility that temporal bias in the collection of some variables could confound their effects with this strong temporal trend. To control for this possibility, we fit the same models using temporal subset of data and without using training end date to confirm the robustness of non-temporal variables (Fig. 8 in Supplementary Materials).

Features of the postdoctoral mentor generally had greater influence on trainee success than the graduate mentor. This suggests a dominant role of postdoctoral advisors on the future career of protégés. The age of the postdoctoral mentor contributed to the likelihood of the protésecuring a permanent position, with older postdoctoral mentors improving the odds. In contrast, graduate mentor age did not have a significant influence.

Network variables were broadly found to have a larger contribution to overall model performance than publication variables (Fig. 5). This relatively greater influence is consistent with the Shapley values and the variable ordering in Table 1, and is found across all formulations of the model studied (Table 3).

We restricted our main analysis to data that was available before the end of training, so as to avoid any confound associated with continuing versus not continuing in academia. To investigate whether the semantic content of papers published after the end of the postdoc continue to influence the career outcomes, we further included them as extra variables in the model. We observed a large explanatory power of late-publication similarity in explaining the continuation in academia, and specially so for the postdoc advisor trainee similarity (supplemental Fig. 10). This strongly suggests that strong ties formed during training and evolving toward a collaboration with the former advisors has a beneficial impact on the trainee career. This also reinforces the idea that the postdoctoral advisor has a larger influence on the future career than the graduate advisor, as was found using variables available at the end of the postdoc (supplemental Figs. 4 and 10).

**Figure 10:**
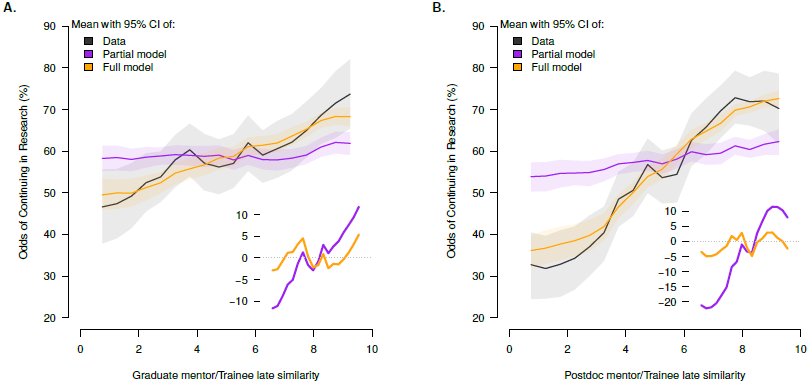
Contribution of publication similarity variables to the odds of continuing an academic career. In each of these plots, the data is shown in black, and the cross-validated predictions of the model including this variable is shown in orange (“full model”). To visualize the contribution of each variable, we also display the cross-validated predictions of the model without this variable in purple (“partial model”). Lines and shaded areas represent respectively the mean values and their 95% bootstrapped confidence interval. Overall, proximity with the postdoctoral advisor has a larger influence than proximity with the graduate advisor. Also, the mentors/trainee proximity displays a larger explanatory power when computed on publication data after the end of training (compared to data available at the end of the postdoc, cf. supplemental Fig. 9), with higher similarity linked to better odds of continuing in research.

### 2.5 Nonlinear influence of mentor graph distance

Mentor graph distance showed a clear inverse relationship with mentor publication similarity (Fig. 2H), which has large explanatory power in the model (Table 1). Indeed, these two variables have a weak but significant correlation (r = −0.192 and p < 0.05, Fig. 6A). Thus, although mentor graph distance had low Shapley value and low importance according to the forward and backward CSA algorithms (Table 1, Fig 6B), we considered the possibility that it might somehow influence trainee outcomes. Mentor graph distribution shows a striking bimodal distribution that suggests a more complex nonlinear relationship with other model variables (Fig. 6C). The distribution of mentor graph distance is broadly similar for trainees who did or did not continue in academia. However for trainees with very short mentor graph distance (< 4 steps) the probability of continuing in academia appears to be consistently lower. We grouped the data into two categories, *tight-knit mentorships* with mentor graph distance less than 4 and *out-of-nest mentorships* with mentor graph greater than or equal to 4. In this case, we do see a different distribution of tight-knit and out-of-nest mentorship groups for the two different outcomes (*p=0.0072*, Pearson’s χ^2^ test using 100,000 Monte Carlo permutations). Thus trainees of advisors that are closely connected in the mentoring graph may be at a disadvantage in acquiring independent research positions, although a larger dataset will confirm that this effect is a confound with other features, in particular the high publication similarity associated with closely related mentors.

## 3. Discussion

We identified a set of factors related to mentorship that influence the academic success of postdoctoral trainees in biomedical research. We considered two measures of academic success: obtaining an independent research position and the number of researchers trained (proliferation rate) once a position is obtained. The main factors influencing the likelihood of obtaining an independent research position were (a) the date of entry in the job market, (b) the success of their mentors (i.e., their proliferation rate), (c) the ability of the trainee to synthesize between the intellectual output of their mentors, and lastly (d) the professional age (time since graduation) of their postdoctoral mentor. The main drivers of the trainee’s proliferation, after obtaining an independent position, were the mentors’ training rates and publishing research that was similar to that of their graduate mentor. Postdoctoral training duration and the professional age of the graduate mentor were found to be irrelevant to trainees success. Below, we discuss these findings and their relevance to the existing literature.

### Trainee synthesis

This study reveals the importance of intellectual synthesis in the pursuit of an independent research career in life sciences. Trainees tended to be more successful if publications by their graduate and postdoctoral had weak semantic similarity, suggesting that their work links ideas and/or approaches from two previously disparate subfields. This finding can be framed within the weak-ties theory of Granovetter ^23^ which emphasizes the importance of bridges between interconnected communities. Applied to academic research, this theory posits some advantages for trainees of mentors belonging to different scientific communities. In their early careers, trainees of disparate mentors benefit from the larger combined network of their advisors, which may be instrumental in creating more professional opportunities and thus helping to secure a permanent position ^22,23^. In addition, researchers bridging disparate scientific communities are in a position to diffuse their innovations more effectively, allowing them to reach more peers and possibly garnering more recognition in their late careers ^43^.

For these beneficial effects to arise, the trainee’s research must be able to act as a bridge. Thus, it is important that the work maintains some similarity with that of the graduate and postdoctoral mentors. Indeed successful trainees tended to have strong semantic similarity to both their graduate and postdoctoral mentors, consistent with a meaningful intellectual impact by both mentors on the trainee’s work. This effect persists and is amplified when considering publications made after the completion of postdoctoral training, suggesting long-lasting benefice of trainees synthesis.

Given that having mentors with dissimilar research positively impacts trainee careers, one might also expect a large mentor network distance to also have a positive influence. However, mentor network distance does not predict trainee continuation or proliferation (Table 1, Fig. 6B). This may reflect the fact that mentor graph distance is too crude a measure of intellectual similarity to provide significant predictive power. Alternatively, there may be a competing social factor, where trainees with closely related mentors may follow a different career trajectory than other trainees. The bimodal distribution in the histogram of mentor graph distance suggests that there are in fact two groups of trainees. Moreover, the decreased odds of having a scientific progeny when co-mentors are linked by three or less mentoring steps supports a hypothesis that the associated lack of diversity could be detrimental to success. Investigation of these fine patterns of close-relationships in training deserve attention in future work.

### Influence of mentors’ success

Another insight of our study is to show that mentors’ training rate positively influences the trainee’s training rate. The link between mentors’ and trainees’ success in academia has been reported previously. In particular, the mentor’s prestige has been shown to be correlated with the trainee’s publication rate^13^. Mentors’ research productivity has also been shown to have a direct impact on both the prestige of the first professional appointment of the trainee, and on research productivity at later stage of their careers ^31^.

This studies reveals a novel finding that the proliferation of the postdoctoral mentor has a greater effect than the graduate mentor on the odds of securing a permanent position (Fig. 4). This trend is consistent with the study of Long et al. ^32^, which showed that the prestige of the postdoctoral institution has a stronger positive effect than that of the doctoral institution for a cohort of biochemists graduated between 1957 and 1963. The greater influence of the postdoctoral advisor is consistent with the idea that the postdoc is a launching pad for an independent career, where scientists build up critical secondary skills (be it network connections, grant funding track-record or technical expertise) required for obtaining a permanent research position.

Overall, our study justifies the long-standing advice that prospective students should look at the training track-record of potential mentors to assess their quality ^5,24^. We show here that besides boosting the odds of securing a permanent research position, highly prolific mentors also tend to have highly prolific trainees, a desirable quality as it is globally linked to academic achievements ^7,14,34^.

### Postdoc duration

Although it is sometimes thought that postdoctoral training duration has increased in recent decades, quantitative reports have consistently showed a stabilization (e.g. ^27^). This previous work draws on data from the Survey of Doctorate Recipients (SDR), which is limited to US-graduated researchers. Thus it discount trends for international postdocs, who often remain longer in postdoctoral positions due to visa limitations ^30^. In the current study, We report that the total duration of postdoctoral training in life science has increased, from less than 3 years in 1970 to 4.1 years in 2015 (Fig. 2F). We also report that the proportion of long postdocs (> 8 years) has remained stable at around 10%. The absence of an increasing trend in long-term positions may be seen as encouraging. Yet, the situations of the 10% “permadocs” remains worrisome and fits in the narrative that sometimes, postdocs are more a source of cheap labor than a meaningful step of career development ^19^.

Previous work on the influence of postdoctoral training time on the odds of securing a permanent position show opposite results. Yang and Webber ^51^ recently showed that chaining 2 to 4 postdoctoral appointments nearly doubled the odds of obtaining an academic position, but did not enhance the long-term productivity of researchers. In a study on biochemists, Nerad and Cerny ^37^ showed, on the contrary, that relatively short postdocs (< 5 years) led to better career outcomes in academia. The objective benefits of completing several postdocs must also be put in balance with a form of survivor’s bias, where those that can afford the cost of long-term postdocs tend to exhibit characteristics correlated with the odds of securing a permanent positions ^51^. In particular, long-term postdoctoral trainees fit more closely to the profiles of tenured researchers, as they tend to originate less often from under-represented minorities and are more often males ^9,46^. The impact of long-term postdoctoral training is then hard to assess in isolation (e.g., women’s more frequent departure from science to take care of children is not linked to scientific ability and rather finds its root in traditional family structures, cf. Chen et al. ^9^, Ginther and Kahn ^21^, Martinez et al. ^35^). Our study contributes to this ongoing debate by showing no systematic link between training duration and academic success. Yet, our study can not rule out the possibility of a spurious negative effect of long cumulative postdoc duration that is masked by independent variables like gender or minority status.

### Mentor experience

The academic age of the postdoctoral mentor at the time of training had a weak but significant positive impact on academic continuation (Figs. 4 and 9F). On the contrary, graduate mentor age was not informative of academic success. Several factors might explain the positive impact of more experienced mentors: greater experience, more material resources in a stable, well-funded laboratory, and better social connections in the network likely to hire the trainee for an independent position. Future studies controlling specifically for these factors are required to determine their relative importance.

Of interest, the study of Malmgren, Ottino and Nunes ^34^ reported opposite effects in the field of mathematics, where the age of the graduate mentor was negatively linked to trainee proliferation. In our study of biosciences, we found a positive effect of postdoctoral mentor academic age on securing a permanent position; and no effect of graduate mentor age on either continuation or proliferation. This discrepancy may arise from the different populations of researchers sampled. This study considered scientists trained by both a graduate and postdoctoral mentor, whereas Malmgren, Ottino and Nunes ^34^ limit their analyses to dyads formed by a graduate advisor and student. In addition, in the field of mathematics, postdoctoral training is much less common, and the prospects of securing a faculty position are twice as good as in other scientific fields ^4^.

## 4.Acknowledgements

This project was funded by a NSF EAGER Award (#1646635) and a Metaknowledge Network Grant to SVD, and AWS Cloud Credits for Research to JFL.

## 5. Methods

### 5.1 Data preparation

The goal of the modeling effort was to assess and predict success in academic research careers. Many different notions of success can be put under this wide umbrella. In this study we chose to quantify trainee proliferation, the number of scientists trained by a scientist, as the measure of academic success ^15,34^. Data for the current study were drawn from the Academic Family Tree, an online, crowdsourced database of mentoring relationships. The database records the identity of the mentor and trainee, the type of relationship (graduate or postdoctoral), and the start and end year of the relationship. Researchers are also linked to publications they have authored that are listed in the Medline database, using a semi-automated algorithm based on string matches to their name and the names of associated trainees and mentors ^52^. Because each trainee can themselves be a mentor for subsequent trainees, the database is represented as a directed graph tracing the growth of academic fields across multiple generations of researchers (Fig. 1A). In order to normalize proliferation measures across mentors and trainees who might still be at different stages of their careers, we computed *proliferation rate*, the average number of trainees per decade since becoming an independent researcher.

As of August 2017, the Academic Family Tree dataset contained 670,000+ researchers across more than 30 fields. Data collection for the Academic Family Tree initially focused only on the field of neuroscience ^15^. Thus this field is more completely represented than fields added more recently. However, its overall properties are similar to other life science fields, including the mentors proliferation rate (Fig. 7). We thus pooled together all fields of life science in this analysis.

### 5.2 Selection criteria

We identified 20,695 triplets in the Academic Tree database, each consisting of a trainee with one graduate and one postdoctoral mentor. In some outlier cases, more than four graduate or postdoctoral advisors were listed, resulting in the same protégés contributing to many triplets. To avoid over-counting these trainees, we removed data for trainees with more than four graduate or postdoctoral mentors. This resulted in 18,856 triplets encompassing 12,853 unique trainees, 9,111 unique graduate mentors and 7,322 unique postdoc mentors.

### 5.3 Missing date inference

Start and end dates of training were entered optionally by users through the web interface. In 49% of the triplets, both start and end dates were available. Training end dates were more relevant to the analyses presented here, as they marked the transition to the status of independent researcher. When both dates were missing, we inferred the end date by identifying the earliest commonly authored publication and adding the field-specific median lag between the start year and first publication. This was performed separately for graduate and postdoctoral training, based on local regression models (LOESS, ^11^) to account for changes in training duration over time. For trainees without an end date but with a start date, we estimated the missing date information by adding the mean difference computed from trainees with complete data, again adjusting for changes in the duration of training periods over time. Using this approach, we were able to assign end dates for training period for 90% of the graduate and 89% of the postdoctoral relationships (see Supplementary Table 2). Overall, 15,583 triplets (83% of the triplets) had dates that could be fully inferred, both at the graduate and postdoctoral levels.

**Table 2:** Statistics of data availability based on publications and manually entered dates

| Availability of date information |  |
| --- | --- |
| Start of PhD | 51% |
| End of PhD | 66% |
| Earliest publication date with graduate advisor | 74% |
| At least one graduate date available | 90% |
| Start of Postdoc | 57% |
| End of Postdoc | 39% |
| Earliest publication date with graduate advisor | 71% |
| At least one postdoc date available | 89% |

**Table 3:** Impact of model types and predictors on predictive accuracy. The mathematical models compared are: the hurdle Poisson model (HP), hurdle negative binomial models (HNB), zero-inflated Poisson models (ZIP) and zero-inflated negative binomial models (ZINB). Predictors were grouped under the same categories as in Table 1: “temporal” for the year of the end of the postdoctoral appointment as well as the total duration of postdoctoral training; “network” for the proliferation of mentors, their academic age at training and their common ancestor distance; and “publications” for the publication similarity between mentors (prior to meeting the trainee) and the publication similarity between postdoctoral trainee and mentor (prior to training). The values displayed are cross-validated log-likelihood aligned on the best model (“ZINB” model using “network + publications + temporal” variables), with higher values denoting more accurate models.

| | $\Delta\mathcal{L}$ in cross-validation | | | |
| --- | --- | --- | --- | --- |
|  | HP | ZIP | HNB | ZINB |
| temporal | −10.24 | −10.23 | −0.47 | −0.45 |
| network | −9.45 | −9.40 | −0.54 | −0.30 |
| publications | −10.42 | −10.38 | −0.71 | −0.52 |
| temporal + network | −9.12 | −9.11 | −0.07 | −0.05 |
| temporal + publications | −10.10 | −10.09 | −0.39 | −0.38 |
| network + publications | −9.31 | −9.26 | −0.47 | −0.24 |
| temporal + network + publications | −8.97 | −8.96 | −0.03 | **0.00** |

### 5.4 Semantic analysis

Academic Family Tree researchers were linked to publications in the Medline and Scopus databases using a simple disambiguation procedure. Publications for which an author name matched a researcher name were attributed to that researcher with high (likely match) or low probability (unlikely) based on clusters in the co-author network ^52^, co-author name matches to adjacent genealogy nodes (i.e., the researcher’s mentor or trainee), and Scopus author identifiers. Website users could then curate publication attributions. As of August 2017, 19.3 million publications have been scanned by the automated system. 197,736 publications for 6,607 researchers have been curated by users, in 90% agreement with the automated system.

For linked publications, we performed latent semantic analysis on abstract text to produce a 400 dimensional vector representing semantic content of each researcher’ publications (cf. Fig. 2B and Achakulvisut et al. ^1^, Deerwester et al. ^16^, Hofmann ^25^, Pedregosa et al. ^40^). Prior to dimensionality reduction, abstracts were pre-processed with stemming, rare word removal, and English stop words removal using the Science Concierge tool suite ^1^. The vector space was then generated by applying a term-frequency inverse document-frequency (TF-IDF) transformation to abstracts for about 90,000 authors, followed by truncated singular value decomposition to the 400 dimensions with greatest variance across authors. Semantic similarity between two researchers was computed by the cosine distance between the vector average across their publications (Fig. 2C). For computing similarity between the two mentors in a triplet, we included only publications prior to any co-authored publication with the trainee. For the predictive model of trainee success, we computed graduate mentor/trainee and postdoc mentor/trainee similarities based only on non-coauthored publications before the end of postdoctoral training. Thus all publication variables were computed from data available before the end of training.

### 5.5 Predictors of success

We hypothesized that several network and semantic variables could predict trainees success:

- *training end year:* year a trainee completed their last postdoctoral fellowship.
- *postdoc duration:* total number of years of postdoctoral training.
- *graduate mentor proliferation, postdoc mentor proliferation*: the average number of the mentor’s graduate and postdoctoral trainees per decade, computed the same way as the trainee proliferation rates.
- *mentor graph distance:* minimum number of steps to pass through a common ancestor between graduate and postdoctoral mentors in the mentorship graph (Fig 1A).
- *graduate mentor age, postdoc mentor age*: number of years since the mentor completed their own training (“academic age” in ^13^).
- *mentor publication similarity:* publication similarity (cosine distance between average publication vectors) between mentors for papers published before they started training the protégés and excluding any co-authored publications.
- *graduate mentor/trainee similarity*, postdoc mentor/trainee similarity: publication similarity between trainee and mentor for publications before the end of postdoctoral training and excluding any coauthored publications.
- *publications with graduate mentor, publications with postdoc mentor:* number of co-authored publications between mentor and trainee prior to the end of postdoctoral training.

To avoid bias in the similarity measure due to co-authored publications (which would have artificially increased publication similarity), we specifically excluded them in the publication similarity computations. That is, graduate mentor/trainee similarity was computed using publications where they do not appear as co-authors. In practice, the publication corpus of the trainee was thus mostly composed of publications co-authored with the postdoctoral advisor.

Data for some variables was only sparsely available. In particular, mentor/mentor and mentor/trainee publication similarity could be computed only for 50% of triplets, as it required identifying publications by each researcher. Likewise, mentor academic age was not widely available, and could be inferred only for 51% of the triplets (see criteria for inference above). Typically, these variables were harder to identify for mentors, as publication data are limited for earlier dates in the Medline and Scopus databases. Thus, of the total number of 18,856 triplets identified in the database, 15,363 had complete temporal information (training end year and postdoc duration), 6,210 had a complete set of mentorship network variables (mentor proliferation, age, and graph distance), and 9,513 had a complete set of semantic variables (mentor/mentor and mentor/trainee similarity and number of co-authored publications). Overall 4,157 triplets satisfied the criterion for a complete data set. From this set of triplets, trainees who completed their training after 2007 were excluded from modeling to take into account the time needed to train students or postdocs when continuing in academia, resulting in a final number of 1,345 triplets used to fit the model.

### 5.6 Model framework

Only a fraction of academically trained individuals go on to have an academic career, and those who do not pursue an independent academic career generally do not have an opportunity to train someone. Thus, overall trainee proliferation depends on two factors: first, whether the postdoctoral researcher secured a permanent position with the opportunity to train new researchers, and second, how many individuals they trained during their subsequent academic career. To account for these two possibilities, we adopted a zero-inflated model formalism. In this framework, the probability of continuing in research is modeled by a binomial variable and the proliferation of researchers that moved on to a permanent research position is modeled by a count variable. Given a vector of predictor variables, *X* (Section 5.5), the model simultaneously describes *π* (*x*), the probability of continuing in an academic career after postdoctoral training, and *f* (*X*), the expected proliferation for those who do continue, as:

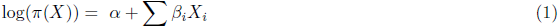

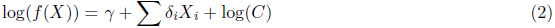

Parameters *β* _i_ and *δ* _i_ indicate the relative weight of the *i* ^th^ variable in predicting *π* and *f*, respectively. The career length is introduced as an offset, log (*C*), because we are ultimately interested in comparing training rates of mentors and *f* (*X*) represents the total count of trainees. Career length is computed as the difference between the year training was completed (end of the last postdoctoral fellowship) and the current year, capped at 45 years (the longest career length reported in the database). Note that in contrast with the usual statistical convention, we define *π* (*X*) as the probability of continuation and not the probability of zero-inflation, which is 1 *-π* (*X*) ^53^.

The interpretation of coefficients differs from standard linear regression due to the presence of the log-link (^6^, ch. 3-4). A change of one unit in the predictor *X _i_* corresponds to multiplying the chance of continuing an academic career by exp(*β _i_*) (log-binomial model) and multiplying the expected trainee proliferation by exp(*δ _i_*) (count model). The exponentiated values of the intercepts, exp(*α*) and exp(*γ*), respectively indicate the baseline continuation probability and proliferation.

In this modeling framework, it is assumed than any researcher who has trained at least one individual has continued in an academic career. However, researchers without a trainee have not necessarily ended their academic career. Such a modeling choice is well-suited to count data with zero-inflation. It is preferred over a simpler linear regression for the following reasons: (a) two distinct processes leading to the absence of trainees are modeled explicitly, (b) correct boundary conditions are enforced by the model design (*i.e.*, the risk of stopping one’s academic career is guaranteed to fall in the range [0,1] and trainee proliferation is never negative) and (c) the number of trainees is not assumed to be normally distributed and can display the over-dispersion expected with count processes.

To confirm that our choice of model formulation and predictors was appropriate, we compared its goodness-of-fit against several alternative formulations (hurdle and zero-inflated, with Poisson and Negative Binomial count models) and predictor sets (Table 3). For each configuration, we evaluated the *predictive* log-likelihood, computed on held-out data that was not used for fitting ^20,45,47^. This cross-validation framework is useful for comparing models that do not assume normally distributed errors and that differ in their number of free parameters ^20^. More specifically, we used *k*-fold cross-validation ^45^, where the data is split into *k* equal-sized random folds, *y* _1_, *…, y* _*k*_. We define *θ* ^*-j*^ as the model parameterization (*β* and *δ* from Eq.1) fit by maximizing log-likelihood on all folds except *y* _*j*_. The predictive log-likelihood in cross-validation *ℒ* is then,

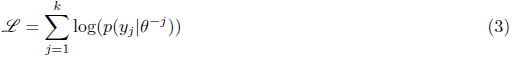

The final estimate of *ℒ* is derived as the average of 100 iterations made each on a different random 10-fold partition. The zero-inflated and hurdle models were optimized using the maximum likelihood procedure implemented in the R package “countreg” ^53^.

### 5.7 Ranking predictors

Shapley values provide an unbiased assessment of the contribution of individual predictor variables to model performance when they are not entirely independent. This metric was originally developed in the field of game theory to score the contribution of each player (here, predictor variable) to coalitions ^3,44^. The Shapley value is computed by considering all possible combinations of predictors and seeing how changing the predictor composition alters model performance (here, cross-validated log-likelihood). Formally, given the log-likelihood *ℒ* and the set of predictors *M*, the Shapley value *ζ*_*i*_ of the predictor *i* is defined as:

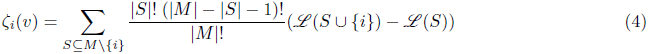

In regression, a notion equivalent to the Shapley values has been developed to quantify the relative importance of regressors by averaging the goodness-of-fit over all possible combinations of variables ^10,29^, in what amounts to a rediscovery of the Shapley values ^48^. Cohen et al. ^12^ proposed an alternative way to use Shapley values in the context of classification, by using it in an iterative variable selection algorithm. Their Contribution-Selection Algorithm (CSA) has a forward and a backward version, which consist in iteratively adding (or removing for the backward version) the variable with the best (or worst) Shapley value.

In this study we computed the cross-validated log-likelihood on the entire set of permutations using all or a subset of the variables. We then considered four ways to select the relevant variables to quantify the odds of continuing in academia and the proliferation rate when continuing in academia, namely: with a brute-force approach (picking the combination maximizing cross-validated log-likelihood); with Shapley value computed on the full set of possible combinations (as in ^10^); and with the forward and backward versions of the CSA (^12^ - see Table 1).

### 5.8 Sensitivity and uncertainty analysis

We computed 95% bootstrapped confidence intervals for descriptive statistics of the dataset and model prediction ^18^. They are shown as error bars and shaded areas throughout the figures.

### 5.9. Data availability

Data from the Academic Family Tree is licensed for re-use with attribution (CC-BY 3.0) and is available through the web portal https://academictree.org or upon request to.

## Supplementary information

### Availability of date information

### Model and variable selection

## 7. Time-dependence of regression coefficients

Fig. 8 shows the regression coefficients obtained when training the model on temporal subsets of the data6 and without the time-controlling variable of “postdoc end year”. Except for this omission, the variables7 included in the model were the ones obtained after the selection process (cf. Table 1 in main text). The8 optimized coefficients from Fig. 8 display much more variability than with the regression shown in main text,9 due to lower sample sizes and the exclusion of temporal variables, but overall display similar trends as the full model.

## 8. Cross-validated model predictions and data

